# Inducible CreER^T2^ mouse lines for characterization of retinal bipolar cell subtypes

**DOI:** 10.64898/2025.12.30.697011

**Authors:** Ebenezer J. Quainoo, Xiaoling Xie, Lotem Kol, Joseph Samson, Lin Gan

## Abstract

Bipolar cells relay visual signals from photoreceptor cells to ganglion cells in the retina. Fifteen bipolar cell subtypes have been identified in the mouse retina. These subtypes are classified as ON and OFF bipolar cells based on their response to light stimulus or rod and cone bipolar cells based on their connection to photoreceptor cells. However, the unique structural and functional role of these subtypes in the processing of visual information is not fully known due to the inadequate tools and models available for their characterization. In this study, to trace the lineage and characterize *Vsx1*, *Lhx3*, and *Lhx4* – expressing bipolar cell subtypes in developing and adult mouse retina, we developed inducible Cre lines under the promoter of *Vsx1*, *Lhx3*, and *Lhx4* and crossed them with ChR2EYFP reporter mice. Lineages of cells expressing *Vsx1*, *Lhx3*, and *Lhx4* after Cre induction in the adult and postnatal mouse retina were then characterized. ChR2EYFP expression driven by *Vsx1*-CreER^T2^ was detected in type 2, 6, and 7 bipolar cells, *Lhx3*-CreER^T2^ in type 1b, 2, and 6 bipolar cells, and *Lhx4*-CreER^T2^ in type 2, 3, 4, and 5 bipolar cells as well as cone photoreceptor cells in the adult mouse retina. These bipolar cell subtype-specific inducible Cre mouse lines serve as efficient tools for elucidation of the mechanisms that control bipolar cell subtype development and function in the retina.

**Significance Statement:** Bipolar cells are central connectors of the outer and inner retina and initiate processing of complex visual information. Bipolar cells differentiate into more than fifteen subtypes during development, and their structural diversity has been well studied. However, the unique contribution of these subtypes to the processing of visual information is poorly understood due to inadequate tools. In this study, we develop and characterize three inducible Cre mouse models for in vivo and temporally defined genetic manipulation in bipolar cell subtypes in the developing and adult mouse retina. These models serve as tools to elucidate the mechanisms that regulate the development and function of bipolar cell subtypes in the mouse retina.

## Introduction

The retina is a light sensitive neural tissue that lies at the back of the mammalian eye. It consists of neuronal cells and supporting glial cells arranged to form a laminar structure. Photoreceptor cells, the modified sensory neurons, receive and convert light signals into electrical signals for processing and transmission through horizontal, bipolar, and amacrine cells, and output retinal ganglion neurons to visual processing centers in the brain (Figure 1A).

**Figure 1.**
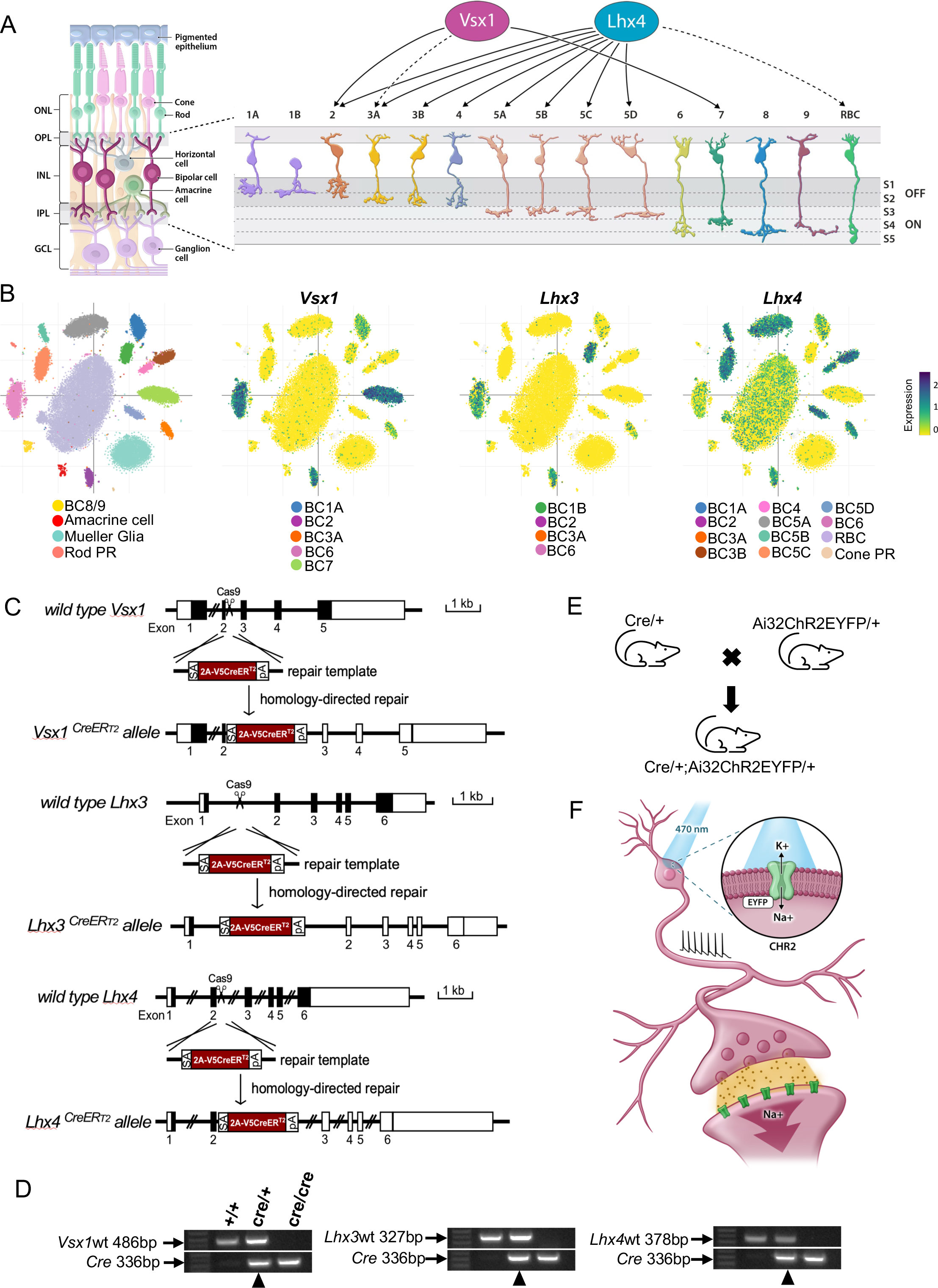
Bipolar cell subtype-specific CreER^T2^ mouse lines. ***A,*** Schematic of retinal cross section highlighting 15 bipolar cell subtypes in the inner nuclear layer (INL); including Type 1 – 4 (OFF bipolar cells) with axon terminal stratification in sublaminae 1 and 2, and Type 5 – 9 as well as rod bipolar cell (ON bipolar cells) with axon terminal stratification in sublaminae 3, 4, and 5 in the inner plexiform layer (IPL). *Vsx1* is expressed in type 2 and 7 bipolar cells and transiently expressed (dashed lines) in type 3a bipolar cells (Shi et al. 2011; 2012). *Lhx4* is also expressed in type 2, 3, 4, and 5 bipolar cells and transiently expressed in rod bipolar cells (Dong et al. 2020). ONL, outer nuclear layer; OPL, outer plexiform layer; GCL, ganglion cell layer; RBC, rod bipolar cell; S, sublaminae ***B,*** Scatter plot of bipolar cell single cell RNA sequencing (Shekhar et al. 2016) showing *Vsx1* expression in type 1a, 2, 3a, 6, and 7 bipolar cells, *Lhx3* expression in type 1b, 2, 3a, and 6 bipolar cells, and *Lhx4* expression in type 1a, 2, 3, 4, 5, 6, and rod bipolar cells as well as cone photoreceptors in P17 mouse retina. ***C,*** Schematic diagram of CreER^T2^ knock-in into *Vsx1*, *Lhx3*, and *Lhx4* utilizing CRISPR-Cas9 and homology-directed repair. CreER^T2^ knock-in allele contains a splicing acceptor site (SA), P2A peptide, V5 tag, and a polyA (PA) for transcription termination. ***D,*** Genotyping of *Vsx1*, *Lhx3*, and *Lhx4* wildtype, heterozygous, and homozygous CreER^T2^ knock-in mice. Heterozygous knock-in mice (black arrowhead) were selected for characterization. ***E,*** Schematic of CreER^T2^ knock-in mice cross to Ai32ChR2EYFP mice to generate double heterozygous CreER^T2^/+; Ai32ChR2EYFP/+ mice. ***F,*** Illustration of light-induced opening of ChR2 with the initiation of an electrical activity in a bipolar interneuron.

Bipolar cells are first order interneurons that connect, process, and relay visual information from the outer retina to the inner retina. Their cell bodies lie in the outer part of the inner nuclear layer with axons and dendrites extending into the outer and inner plexiform layers of the retina. Bipolar cells, as other neuronal cell types in the retina, comprise many subtypes. These subtypes are broadly categorized into ON and OFF bipolar cells depending on their response to light stimulus or rod and cone bipolar cells depending on the type of photoreceptor cell inputs they receive (Hack, et al. 2001; Haverkamp, et al. 2000; Masu, et al. 1995; Tsukamoto, et al. 2001). Ghosh, et al. (2004) further classified ON and OFF bipolar cells based on the level of their axon terminal stratification (Figure 1A). OFF bipolar cells axons stratify in sublaminae 1 and 2, and ON bipolar cells axons stratify in sublaminae 3, 4, and 5 in the inner plexiform layer of the retina. Bipolar cells have also been classified based on their gene expression profile; revealing 15 distinct subtypes that correspond to their morphological classification (Shekhar, et al. 2016). Notably, neuronal subtype classification based on their gene expression profile give significant insights into the gene regulatory mechanisms that control subtype formation, differentiation, and function.

All 15 bipolar cell subtypes identified make distinct connections to other neuronal cell types in the retina and forms the initial processing center of the visual system; that is, begins the processing of complex visual information as it travels to visual centers in the brain. As intriguing as bipolar cells are, it is still not clear the factors that control their subtype formation. It is also not known whether these subtypes are functionally distinct, that is, whether each subtype or group of subtypes has specific function in processing of visual information in the retina. These questions warrant the development of tools, models, and techniques for further study of bipolar cells development and function in the retina.

During retinogenesis, homeobox genes encode homeodomain-containing transcription factors that carry out essential roles in retinal progenitor cells specification and cell type differentiation in the retina (Zagozewski, et al. 2014). Among these homeobox genes are VSX1, LHX3, and LHX4 transcription factors that specifically regulate bipolar cell subtype development in the retina (Quainoo and Gan 2025; West and Cepko 2022). Previous studies have characterized the expression of these transcription factors in the retina. *Vsx1*, for instance, has been shown to be expressed in type 7 bipolar cells and OFF bipolar cell subtypes that co-express type 2 bipolar cell markers in adult GUS8.4GFP and *Vsx1*:^t^lacZ transgenic reporter mice (Shi, et al. 2012; Shi, et al. 2011). Transient expression of *Vsx1* is also detected in type 3a OFF bipolar cells within the first two postnatal weeks in the mouse retina (Shi, et al. 2012). *Lhx4* expression, on the other hand, has been reported in OFF type 2, 3, and 4 bipolar cells and ON type 5 bipolar cells with transient expression in rod bipolar cells within the first two postnatal weeks in the mouse retina (Dong, et al. 2019; Dong, et al. 2020) (Figure 1A). A study by Shekhar, et al. (2016) also characterized the gene expression profile of retinal bipolar subtypes in P17 mouse retinas and showed *Vsx1*, *Lhx3*, and *Lhx4* expression in both ON and OFF bipolar cell subtypes. Single cell RNA sequencing data of this study shows *Vsx1* expression in type 2, 3a, and 7 bipolar cells and also shows its expression in type 1a and 6 bipolar cells. *Lhx4* expression, consistent with Dong, et al. (2020), is also detected in type 2, 3, 4, and 5 bipolar cells and *Lhx3* expression in types 1b, 2, and 6 bipolar cells. Weak expression of *Lhx3* is also detected in type 3a bipolar cells, and *Lhx4* in type 1a, type 6, and rod bipolar cells (Figure 1B). These gene expression patterns display the overlapping and distinct expression of *Vsx1*, *Lhx3*, and *Lhx4* in different bipolar cell subtypes during bipolar cell development.

Transgenic Cre reporter mouse models including *Kcng4*-cre, *Neto1*-cre, *Fezf1*-cre, and *Pcp2*-cre have been used to characterize retinal bipolar cell subtypes (Daigle, et al. 2018; Duan, et al. 2014; Zhang, et al. 2004). However, these reporter mouse models are either not inducible or specific to retinal bipolar cells. In this study, we developed and characterized three inducible Cre mouse models using promoters of key transcription factor genes involved in bipolar cell subtype development namely; *Vsx1*, *Lhx3*, and *Lhx4*. Through these models, we also showed the differential transcription factor expression patterns that occur and influence bipolar cell subtype formation during their development in the postnatal retina and in the adult mouse retina. These models allow control of time and cell density during genetic manipulation in bipolar cells and thus provide accessible means and approach for further characterization of mechanisms regulating bipolar cell subtypes specification and function.

## Materials and Methods

### Generation of Mouse Lines

Mice were generated by [Author University] Genome Editing Core using CRISPR/Cas9 approach. Briefly, the DNA repair template containing the CreER^T2^ mutations was synthesized and purified by Integrated DNA Technologies (Coralville, Iowa 52241 USA). Single guide RNA (sgRNA) targeting *Vsx1*, *Lhx3*, and *Lhx4* gene locus was synthesized and purified by Synthego (Redwood City, CA). RNP complex of 60 pmol Cas9 protein (IDT, Alt-R^TM^ S.p. Cas9 Nuclease V3, stock #1081059) and 60 pmol sgRNA was formed and mixed with 300 pmol repair template DNA in 50 ml injection buffer and injected into fertilized eggs from C57BL/6J mice (Jackson Laboratory, stock #000664). Viable two-cell stage embryos were transferred to pseudo-pregnant Swiss Webster females (Taconic Biosciences, Inc., Germantown, NY, stock #SW-F-EF) to generate founder mice. The positively targeted founder mice were identified by external long-range PCRs and Sanger sequencing by Azenta Life Sciences (Research Triangle Park, NC) and were subsequently bred with wild type C57BL/6J mice for germline transmission to generate F1 mice. F1 heterozygous pups with desired mutation were further confirmed by external long-range PCRs and Sanger sequencing. Mice were housed in a standard 12 h light/12 h dark cycle environment and all animal procedures were performed in accordance with IACUC protocols at [Author University] following guidelines described in the US National Institutes of Health Guide for the Care and Use of Laboratory Animals. All mice were crossed to an inducible Cre recombinase-dependent reporter mouse line that expresses ChR2EYFP from the *Rosa26* locus (Ai32, JAX Strain#: 012569). Mice of either sex were used in all experiments.

### Tamoxifen treatment

Three doses of 75 mg/kg tamoxifen were administered through intraperitoneal (IP) injection in adult *Lhx3-*CreER^T2^ and *Lhx4-*CreER^T2^ mice between P50 – P55 and retinal tissues or eyecups were harvested a week after the last dosage. Tamoxifen injection in *Vsx1-*CreER^T2^ mice was administered earlier at P40 for better Cre recombination and reporter expression. For Cre induction in postnatal mice, 50 μl of 20 mg/ml tamoxifen was administered to pups through IP injection at P3, P5, and P7 and eyecups of mice were harvested at P60.

### Immunohistochemistry

Mice were euthanized with CO_2_ and cervical dislocation. Eyeballs were enucleated and briefly fixed in ice-cold 4% PFA for 5 min and cornea, iris, and lens were removed in 1X PBS. Eyecups were fixed in 4% PFA at 4°C overnight and thoroughly washed for 2X 15 min each in 1X PBS. Washed eyecups were then dehydrated with sucrose gradient and embedded and quickly frozen in OCT for cryosectioning. Immunofluorescence staining was performed as previously described (Guo, et al. 2023). Briefly, 14 mm thick cryosections were washed in 1X PBS (pH 8.0), followed by permeabilization in 0.3% PBST (Triton X-100 in 1X PBS). Sections were then blocked with 10% normal horse serum in 0.3% PBST (blocking solution) in a humidity chamber for 1 hr at room temperature. Primary antibodies diluted in blocking solution were then applied on sections and incubated at 4°C overnight. Sections were washed 2X 15 min each in 0.1% PBST after primary antibody incubation and secondary antibodies diluted in blocking solution were applied and incubated for 1 hr at room temperature. Conjugated second primary antibody was applied for overnight incubation at 4°C after washing off secondary antibody in 0.1% PBST. Sections were then washed in 0.1% PBST and 1X PBS after second primary antibody incubation and incubated in DAPI for nuclei staining for 10 min followed by mounting with coverslips after brief washing of DAPI. For wholemount retinas, enucleated eyeballs were fixed in 4% PFA overnight at 4°C. Whole retinas were dissected and fixed in 4% PFA for 1-2 hrs at 4°C and then washed in 1X PBS 2X 15 min each. Immunolabeling was then performed as above, and retina was flat mounted on glass slides for imaging. Primary antibodies used were rabbit anti-GFP alexa fluor 488 (1:500, Molecular Probes #A-21311), sheep anti-VSX2 (1:200, Exalpha #X1180P), sheep anti-calretinin (1:500, Molecular Probes #PA5-95651), goat anti-BHLHB5 (1:1,000, Santa Cruz #sc-6045), mouse anti-calsenilin (1:250, Millipore #05-756), mouse anti-PKARIIb (1:500, BD Biosciences #610625), rabbit anti-opsin red/green (1:150, Millipore #ab5405), rabbit anti-PKCA (1:8,000, Sigma #P-4334). Corresponding secondary antibodies were applied at a dilution of 1:1,000 and are as follows: Alexa-647-conjugated donkey anti-mouse (ThermoFisher Scientific #A-32787), Alexa-647-conjugated donkey anti-goat (ThermoFisher Scientific #A-32849), Alexa-568-conjugated donkey anti-goat (ThermoFisher Scientific #A-11057), Alexa-647-conjugated donkey anti-rabbit (ThermoFisher Scientific #A-31573), Alexa-647-conjugated donkey anti-sheep (ThermoFisher Scientific #A-21448), Alexa-568-conjugated donkey anti-sheep (ThermoFisher Scientific #A-21099).

### Imaging

Retinal sections were imaged using a Leica Stellaris Confocal microscope (Leica Microsystems Inc, IL) with a 20X or 40X oil objective lenses in the 405, 488, 568, and 647 nm laser channels. Z-stacks were obtained whenever appropriate at 0.1-0.5 mm step size for 2D maximum projection of image sections. Channels were overlaid whenever appropriate, and brightness and contrast adjustments were made using Adobe Photoshop (Adobe Inc, San Jose, CA).

### Statistics

Descriptive statistics was employed to determine the distribution and density of bipolar cells in retina sections and wholemounts. Data was analyzed and presented as mean ± SEM using Prism software (GraphPad, San Diego, CA).

## Results

### Bipolar cell subtype-specific inducible Cre mouse lines

To generate inducible Cre knock-in mouse models at *Vsx1*, *Lhx3*, and *Lhx4* gene loci, a gene cassette containing Adenovirus major late intron splicing acceptor sequence, P2A-CreER^T2^ and rabbit beta-globin polyadenylation signal (rBG polyA) was knocked in between exon 2 and 3 of *Vsx1* and *Lhx4* and between exon 1 and 2 of *Lhx3*, for CreER^T2^ expression and rBG polyA transcription termination and gene knock out (Figure 1C, D). Whereas *Vsx1*CreER^T2^ homozygous knockout mice survive postnatally and throughout adult, *Lhx3*CreER^T2^ and *Lhx4*CreER^T2^ homozygous knockout mice were either embryonic or perinatally lethal. Adult heterozygous mice of each Cre knock-in line were therefore used for experiments. These mice were crossed with a Cre-dependent channelrhodopsin-2 fused with enhanced yellow fluorescent protein (ChR2EYFP) reporter mice to generate CreER^T2^;Ai32ChR2EYFP double heterozygous mice for characterization of the expression of *Vsx1*, *Lhx3*, and *Lhx4* in retinal bipolar cell subtypes in developing and adult mouse retinas (Figure 1E). ChR2 is a light-gated ion channel and integral membrane protein that mediates light-induced electrical activity particularly in neurons (Figure 1F). The expression of ChR2EYFP in bipolar cell subtypes optimizes visualization of cell morphology and makes manipulation of light-induced neuronal activity in bipolar cell subtypes feasible (Madisen, et al. 2012).

### Lineage tracing of *Vsx1*, *Lhx3*, and *Lhx4* – expressing bipolar cell subtypes in the adult mouse retina

Cell lineage tracing using inducible Cre models allow temporal control of genetic labeling of specific cells or tissues throughout their development (Kretzschmar and Watt 2012). Thus, to identify bipolar cell subtypes that express *Vsx1*, *Lhx3*, and *Lhx4* in the adult mouse retina, we induced Cre/loxP recombination in adult CreER^T2^;Ai32ChR2EYFP mice and performed immunolabeling on retinal tissues to assess ChR2EYFP reporter expression.

Flat mount immunolabeling showed the expression of ChR2EYFP in the membranes of bipolar cells in adult *Vsx1*, *Lhx3*, and *Lhx4*-CreER^T2^ mouse retinas (Figure 2A). *Lhx4*CreER^T2^ mouse retinas showed a high density of ChR2EYFP-expressing bipolar cells (142.3±3.14 cells/100 mm^2^) compared to that of *Lhx3*CreER^T2^ (27.5±0.62 cells/100 mm^2^) and *Vsx1*CreER^T2^ (24.0±1.11 cells/100 mm^2^) given same dosage and concentration of tamoxifen (3X, 75 mg/kg). This likely correlates to the number of bipolar cell subtypes expressing these genes in the adult mouse retina during the time of Cre induction. In retinal cross-sections, cell bodies of CHR2EYFP-expressing bipolar cell subtypes colocalized with the pan bipolar cell marker CHX10/VSX2 (Burmeister, et al. 1996) in the outer segment of the inner nuclear layer (Figure 2B). Co-immunolabeling of EYFP-expressing bipolar cells with calretinin also showed axon terminal of bipolar cell subtypes laminating in both ON and OFF sublamina layers in the inner plexiform layer (Ghosh, et al. 2004) indicating *Vsx1*, *Lhx3*, and *Lhx4* expression in both ON and OFF cone bipolar cells (Figure 2C). Interestingly, *Lhx4* expression was also detected in subsets of cone photoreceptor cells that co-immunolabeled with red/green opsin (Figure 2D,E).

**Figure 2.**
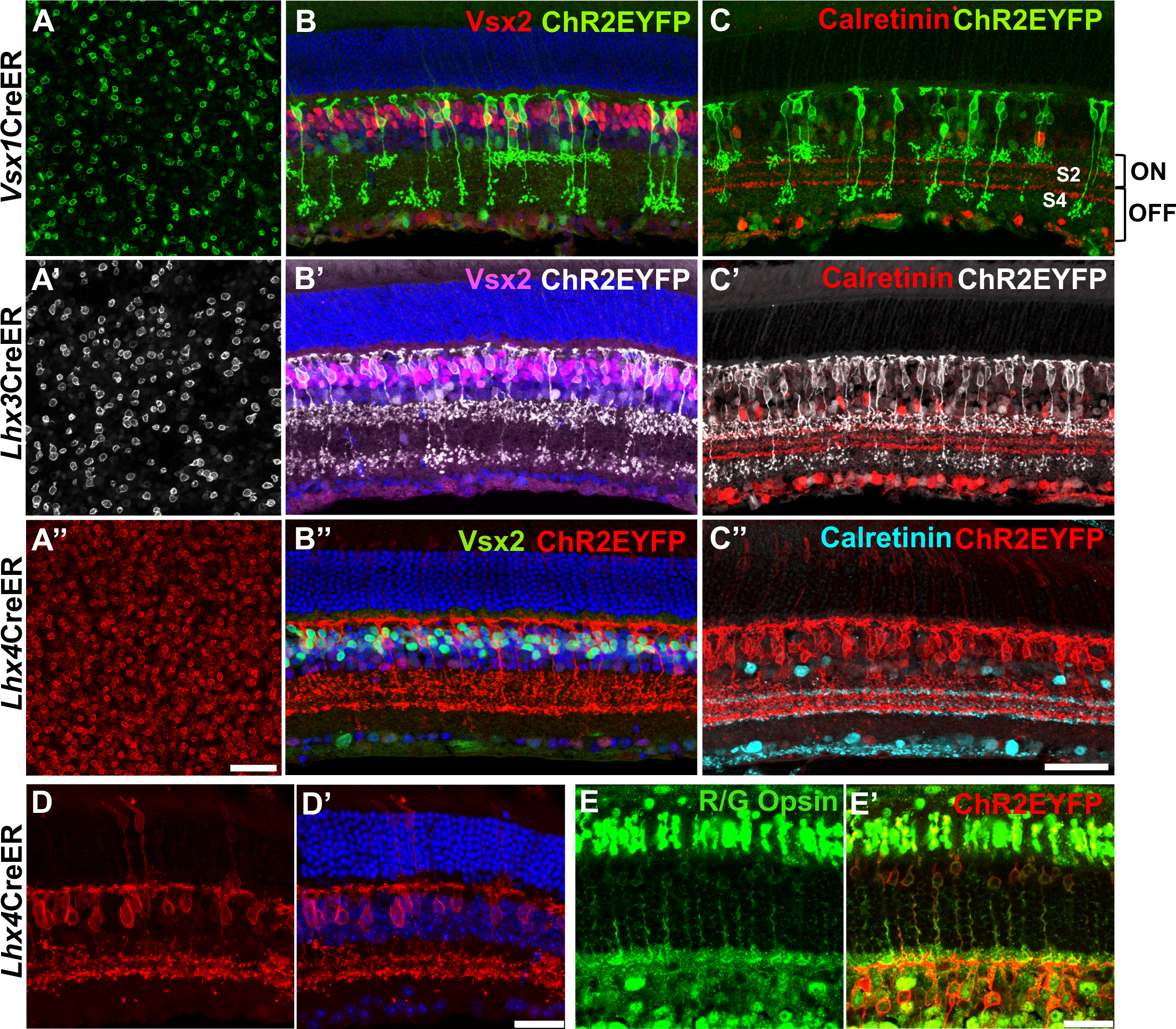
Lineage tracing of *Vsx1*, *Lhx3*, and *Lhx4* in ON and OFF bipolar cells and cone photoreceptor cells in the adult mouse retina. ***A,*** Flatmount retinas showing ChR2EYFP expression in the cell membrane of *Vsx1*, *Lhx3*, and *Lhx4* – expressing retinal bipolar cell somas after 3X tamoxifen dosage. Scale bar in A’’ = 25 μm. ***B,*** Retina cross-sections showing co-localization of *Vsx1*, *Lhx3*, and *Lhx4* – expressing bipolar cells with pan bipolar cell marker Vsx2. ***C,*** *Vsx1*, *Lhx3*, and *Lhx4* – expressing bipolar cells axons stratify in ON and OFF sublaminae layers in the IPL of the mouse retina. Scale bar (retina cross-sections) in C’’ = 50 μm. ***D,*** Maximum projection retina cross-section images showing ChR2EYFP expression in *Lhx4*-expressing bipolar cells and photoreceptor cells. ***E,*** Co-immunolabeling of *Lhx4*-expressing photoreceptor cells with red/green (R/G) opsin. Scale bar (***D***,***E***) = 25 μm.

The wide separation of ON and OFF bipolar cells axon terminal stratification in *Vsx1* and *Lhx3* – expressing bipolar cells made it possible to determine the percentage of ON bipolar cells in *Vsx1*CreER^T2^ (59.9%±1.77%) and *Lhx3*CreER^T2^ (45.3%±4.51%) adult mouse retinas. Using the type 2 bipolar cell marker BHLHB5 (Feng, et al. 2006), we also found that 12.7%±4% EYFP-expressing bipolar cells in *Vsx1*CreER^T2^, 36.4%±3.48% EYFP-expressing bipolar cells in *Lhx3*CreER^T2^, and 19.7%±1.36% EYFP-expressing bipolar cells in *Lhx*4CreER^T2^ adult mouse retinas were type 2 bipolar cells (Figure 3A, B, C). Given about 40% *Vsx1*-expressing OFF bipolar cells – with axon terminal stratification in S1 – and approximately 13% *Vsx1*-expressing type 2 OFF bipolar cells in the adult mouse retina, there remains about 27% non-type 2 OFF bipolar cells (type 1) present in the *Vsx1*CreER^T2^ adult mouse retina. However, a much higher number of BHLHB5+ type 2 bipolar cells (40.3%±2.86%) were detected in *Vsx1*CreER^T2^ retinas when tamoxifen was injected at an earlier timepoint (P7) (Figure 3A’) suggesting *Vsx1* down-regulation in type 2 bipolar cells in the adult mouse retina. Subsets of *Lhx4*-expressing bipolar cells also co-immunolabeled with type 3b bipolar cell marker PRKAR2B (Mataruga, et al. 2007) (23.1%±1.99%) and type 4 bipolar cell marker calsenilin (Haverkamp, et al. 2008) (22.9%±1.19%) indicating *Lhx4* expression in these subtypes (Figure 3D, E).

**Figure 3.**
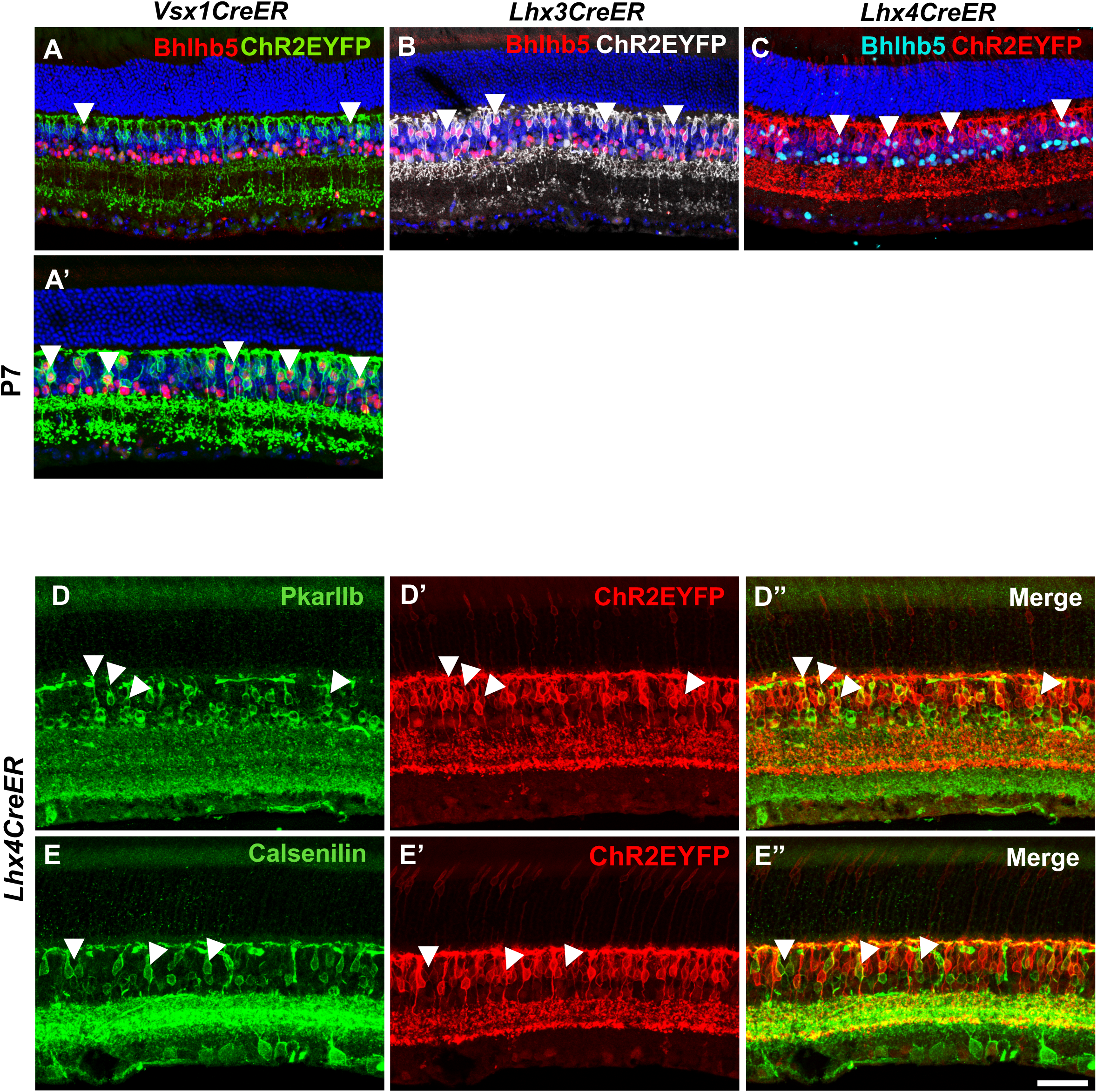
*Vsx1*, *Lhx3*, and *Lhx4* – expressing bipolar cells colocalizes with OFF bipolar cell markers in the adult mouse retina. ***A-C,*** Retina cross sections showing colocalization of type 2 bipolar cell marker Bhlhb5 in subsets of *(A) Vsx1*, *(B) Lhx3*, and *(C) Lhx4*-expressing bipolar cells. ***D*** and ***E,*** Co-immunolabeling of type 3 bipolar cell marker PkarIIb *(D)* and type 4 bipolar cell marker Calsenilin *(E)* with subsets of *Lhx4*-expressing bipolar cells. Scale bar = 50 μm.

Taken together, ChR2EYFP reporter expression patterns show that *Vsx1*, *Lhx3*, and *Lhx4* – expressing cells give rise to both ON and OFF cone bipolar cells and cone photoreceptor cells in the adult mouse retina.

### Sparse labeling and morphology of *Vsx1*, *Lhx3*, and *Lhx4* – expressing bipolar cell subtypes in the adult mouse retina

Sparse labeling is an essential method for visualizing and studying the morphology of individual cells. In an inducible Cre model, this method is achieved by varying and fine-tuning the concentration of Cre-activating ligand. Thus, to sparse label and further characterize *Vsx1*, *Lhx3*, and *Lhx4* – expressing bipolar cell subtypes, tamoxifen concentration for Cre recombination was decreased and varied among *Vsx1*, *Lhx3*, and *Lhx4*-Cre mice to the level where reporter expression occurred in only few interspersed bipolar cell subtypes in the adult retina.

Flat mount staining of sparsely labeled *Vsx1*, *Lhx3*, and *Lhx4* – expressing bipolar cell subtypes showed relatively reduced density of bipolar cells (Figure 5). *Vsx1*-expressing bipolar cell subtypes was sparsely labeled with a single dose of 75 mg/kg tamoxifen and high magnification confocal imaging of sparsely labeled bipolar cells in retinal sections showed *Vsx1* expression in type 2, 6, and 7 bipolar cells (Figure 4A). Axon terminal of type 2 bipolar cells stratified in S1 in the inner plexiform layer and were brush-like with dense varicosities (Ghosh, et al. 2004). Both type 6 and type 7 *Vsx1*-expressing bipolar cells axon terminal stratified in S4. However, type 7 bipolar cells had a more laminated and widespread terminal branching than type 6. Type 6 bipolar cells also tended to have a more elongated cell body than the relatively rounded cell body of type 7 bipolar cells.

**Figure 4.**
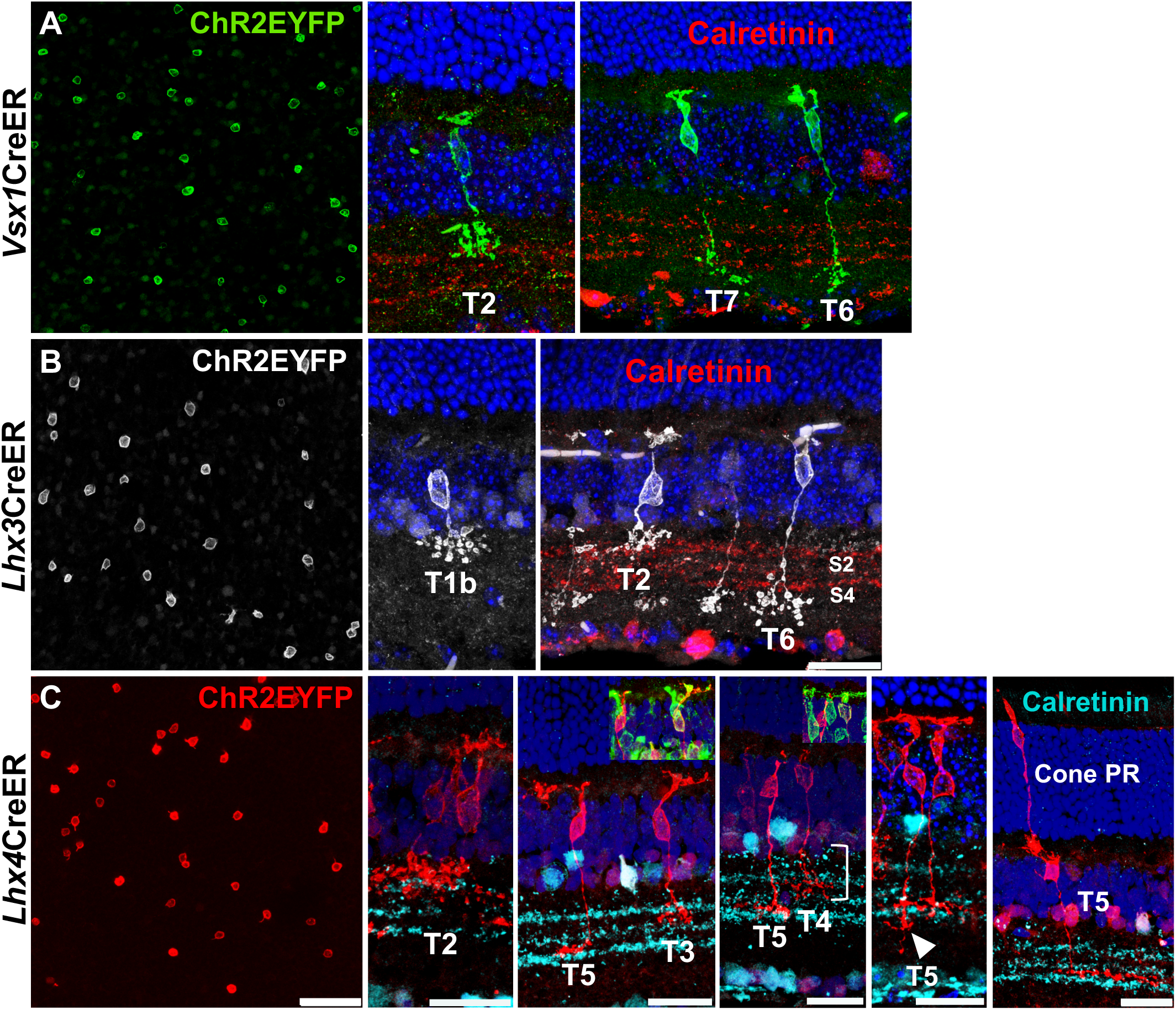
Sparse labeling of *Vsx1*, *Lhx3*, and *Lhx4* – expressing bipolar cells in the adult mouse retina. ***A,*** Sparsely labeled *Vsx1*-expressing bipolar cells in retina flatmount and cross-sections showing *Vsx1* expression in type 2 OFF and type 6 and 7 ON bipolar cells. ***B,*** Sparsely labeled *Lhx3*-expressing bipolar cells in retina flatmount and cross-sections showing *Lhx3* expression in type 1b and 2 OFF and type 6 ON bipolar cells. ***C,*** Sparsely labeled *Lhx4*-expressing bipolar cells in retina flatmount and cross-sections showing *Lhx4* expression in type 2, 3, and 4 OFF and type 5 ON bipolar cells and in a cone photoreceptor cell. Axon terminal stratification of some type 5 bipolar cells extends into sublaminae 4 (white arrowhead). Type 4 bipolar cells have a characteristic large axon arbor thickness (right bracket). Insets show co-labeling of type 3 bipolar cell with PkarIIb and type 4 bipolar cell with calsenilin. Scale bar = 25 μm.

**Figure 5.**
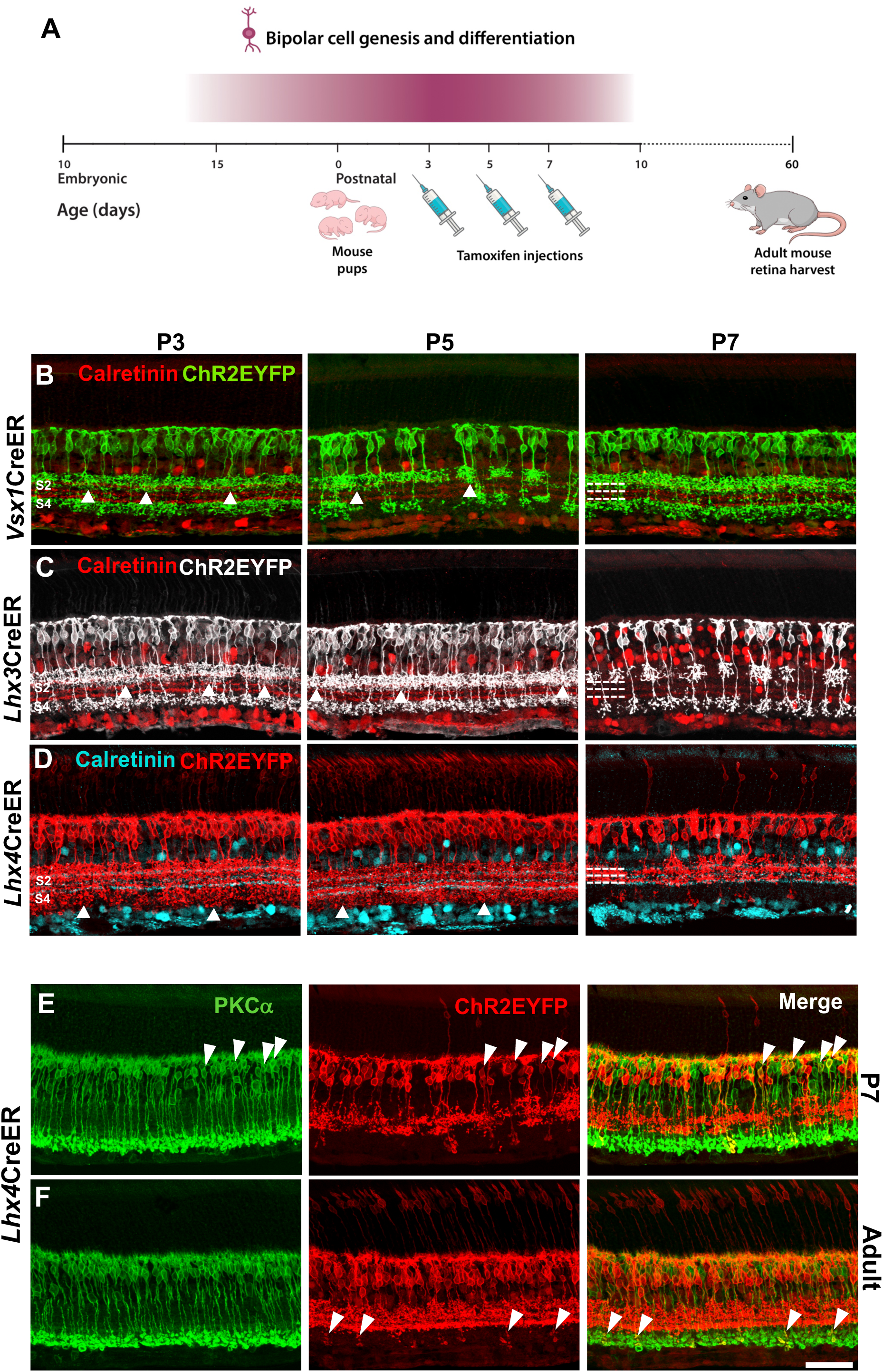
*Vsx1, Lhx3,* and *Lhx4* expression pattern during bipolar cell genesis and differentiation. ***A,*** Schematic of experimental procedure involving tamoxifen injection in P3, P5, and P7 pups and retinal harvest for EYFP expression characterization at P60. ***B,*** *Vsx1*-expressing bipolar cells in P3, P5, and P7 tamoxifen injection mice with axon terminal stratification in S1, S2, and S4 layers in the adult mouse retina. ***C,*** *Lhx3*-expressing bipolar cells in P3, P5, and P7 tamoxifen injection mice with axon terminal stratification in S1, S2 (P3, P5), and S4 layers in the adult mouse retina.. ***D,*** *Lhx4*-expressing bipolar cells in P3, P5, and P7 tamoxifen injection mice. *Lhx4* is expressed in bipolar cells with axon terminal stratification beyond S3 in P3 and P5 tamoxifen injection mice but mostly in bipolar cells with axon terminal stratification in S1, S2, and S3 in P7 tamoxifen injection mice. ***E,*** Co-immunolabeling of rod bipolar cell marker PKCa and subset of *Lhx4*-expressing bipolar cells in P7 tamoxifen injection mouse retina. ***F,*** Co-immunolabeling of rod bipolar cell marker PKCa and *Lhx4*-expressing axon terminals in the adult mouse retina. Scale bar = 50 μm

*Lhx3*-expressing bipolar cell subtypes were sparsely labeled with a single dose of 40 mg/kg tamoxifen. High magnification confocal imaging of *Lhx3*-expressing bipolar cells in retinal sections showed *Lhx3* expression in type 2 and type 6 bipolar cells (Figure 4B). *Lhx3* was also expressed in type 1b bipolar cells and the expression of ChR2EYFP revealed its unique amacrine cell-like cell body-axon unipolar morphology as described by Shekhar, et al. (2016) and Della Santina, et al. (2016).

*Lhx4*CreERT^2^ mice required much lower concentration of tamoxifen – a single dose 15 mg/kg – to sparsely label *Lhx4*-expressing bipolar cell subtypes, highlighting the high density of *Lhx4*-expressing bipolar cells in the adult mouse retina. Lowering concentration of tamoxifen even further (10 mg/kg) only labeled subtypes that had their axon terminals stratifying in S2 (type 3 and type 4). Morphology and axon terminal stratification of sparsely labeled *Lhx4*-expressing subtypes, however, showed *Lhx4* expression in type *2*, 3, 4, and 5 bipolar cells (Figure 4C). Interestingly, type 2 bipolar cells were rarely detected in sparsely labeled *Lhx4*-expressing subtypes suggesting relatively low expression of *Lhx4* in type 2 bipolar cells in the adult mouse retina. Type 3 and type 5 bipolar cells varied in their morphology. However, all their morphological variants had axon terminals stratifying in S2 and S3 respectively (Ghosh, et al. 2004). Some type 5 bipolar cells had their axon terminals extending into S4 and axon terminals of some type 3 bipolar cell variants began their branching in S1. However, type 4 bipolar cells characteristically had a much larger arbor thickness; that is, a higher axon terminal branching depth extending S1 and S2 (Ghosh, et al. 2004; Tsukamoto and Omi 2014). Since axon terminals of both type 3 and type 4 stratified at S2, the most distinctive way to identify them was their co-immunolabeling with subtype-specific markers PkarIIb and Calsenilin (Haverkamp, et al. 2008; Mataruga, et al. 2007). Sparse labeling in *Lhx4*CreERT^2^ retinas also showed cone photoreceptors making synaptic contacts with cone bipolar cells in the outer plexiform layer (Figure 4C).

### *Vsx1*, *Lhx3*, and *Lhx4* expression patterns during retinal bipolar cell genesis and differentiation

Peak genesis of bipolar cells in the mouse retina occurs within the first postnatal week alongside Müller glia and rod photoreceptors (Young 1985). Expression of transcription factors in the retina during this time regulate both the specification, differentiation and subtype formation of bipolar cells (Quainoo and Gan 2025; West and Cepko 2022). The subtype-specific expression pattern of transcription factors in bipolar cells during this time of peak genesis is, however, not clear. To investigate this, we characterized *Vsx1, Lhx3,* and *Lhx4* expression in the retina during the first postnatal week. A single dosage of tamoxifen was injected intraperitoneally on P3, P5, and P7 in *Vsx1*CreERT^2^, *Lhx3*CreERT^2^, and *Lhx4*CreERT^2^ mice and EYFP expression was characterized in the adult P60 mouse retina (Figure 5A). Complete degradation of tamoxifen in mice occurs in about 4-6 days post injection (Valny, et al. 2016). Hence, subtype-specific expression pattern of *Vsx1, Lhx3,* and *Lhx4* in bipolar cells on the day of injection and about 4-6 days post-injection was observed in the P60 adult mouse retina.

Two possible outcomes were expected; that these transcription factors at the onset of bipolar cells specification are expressed in specific bipolar cell subtypes as observed in the adult retina or are turned on and off in different bipolar cell subtypes at different timepoints during bipolar cells development.

*Vsx1* expression in the retina of P3, P5, and P7 tamoxifen injection mice were restricted in ON and OFF bipolar cells that had axon terminal stratification largely in S1 and S4 in the inner plexiform layer (Figure 5B). However, in contrast to *Vsx1*–expressing bipolar cells in adults, *Vsx1* was also expressed in bipolar cells that had axon terminal stratifications in S2 in the P3, P5, and P7 tamoxifen injection retinas suggesting *Vsx1* transient expression in type 3 or 4 bipolar cells during the first postnatal week. Similar observations were also found in the *Lhx3*CreER^T2^ retinas. In contrast to *Lhx3* expression in the adult mouse retina, *Lhx3* expression in P3 and P5 tamoxifen injection retinas were detected in bipolar cells that had axon terminal stratification on S2. *Lhx3*–expressing bipolar cells in the P7 tamoxifen injection retinas, however, showed similar axon terminal stratifications as those observed in the adult mouse (Figure 5C).

As in adults, *Lhx4* was expressed in both photoreceptor cells and bipolar cells during the first postnatal week (Figure 5D). However, unlike in adults, *Lhx4* was expressed in bipolar cells that had axon terminal stratification beyond S3 in *Lhx4*CreERT^2^ retinas injected with tamoxifen on P3 and P5. This shows that *Lhx4* is expressed in many atypical ON cone and perhaps rod bipolar cells during the first postnatal week. In the P7 tamoxifen injection retinas, *Lhx4* expression was mainly restricted to bipolar cells that had axon terminal stratification in S1, S2, and S3 as in adults. However, subsets of *Lhx4*–expressing bipolar cells in P7 tamoxifen injection retinas co-immunolabeled with the rod bipolar cell marker PRKCA (Haverkamp and Wässle 2000) (Figure 5E) indicating *Lhx4* expression in rod bipolar cells during the first postnatal week. In the adult mouse retina, *Lhx4* expression was detected only in the axon terminals of a few rod bipolar cells and suggested *Lhx4* decreased expression or shutdown in rod bipolar cells during the course of development (Figure 5F).

Taken together, *Vsx1*, *Lhx3,* and *Lhx4* shows differential expression in bipolar cell subtypes at different timepoints during bipolar cell development. This shows that differential expression of transcription factors during early bipolar cell development likely regulates bipolar cell subtype formation and maintenance.

## Discussion

Study characterizes the efficiency of inducible CreER^T2^ mouse models and traces the lineage of *Vsx1, Lhx3,* and *Lhx4* in bipolar cell subtypes during their development in the postnatal retina and in the adult mouse retina. Main findings of study show that *Vsx1* – expressing bipolar cells give rise to type 2 and 7 as well as type 6 bipolar cell subtypes, *Lhx3* – expressing bipolar cells give rise to type 1b, 2, and 6 bipolar cell subtypes, and *Lhx4* – expressing bipolar cells give rise to type 2, 3, 4, and 5 bipolar cell subtypes as well as cone photoreceptor cells. *Lhx4* is also shown to be widely expressed in many different bipolar cell subtypes during bipolar cell specification and differentiation in the first postnatal week and expressed in fewer bipolar cell subtypes later during development and in the adult retina. Similarly, *Vsx1 and Lhx3* is shown to exhibit variation in their expression pattern in bipolar cell subtypes during their development in the postnatal retina and in the adult mouse retina.

These inducible CreER^T2^ mouse models are essential tools for studying bipolar cell subtypes development and function and augment the inadequate subtype-specific genetic tools available for the study of bipolar cells. Previous work has shown expression of *Vsx1*, *Lhx3*, and *Lhx4* in bipolar cell subtypes and support the expression pattern of these genes in our study. *Vsx1* has been shown to be expressed in type 2, 6, and 7 bipolar cells with temporal or weak expression in types 1a and 3a bipolar cells (Shekhar, et al. 2016; Shi, et al. 2012; Shi, et al. 2011). Knockout of *Vsx1* results in incomplete terminal differentiation and function of OFF cone bipolar cells and type 7 bipolar cells (Chow, et al. 2004; Shi, et al. 2011). However, the function of *Vsx1* in type 6 bipolar cells in the retina have not been well characterized. Our study shows *Vsx1* expression in OFF bipolar cells and in type 6 and 7 ON bipolar cells in the *Vsx1*CreER^T2^ mouse retina and thus, provides a tool for structural and functional characterization of type 6 bipolar cells. The presence of BHLHB5 – OFF bipolar cells in the *Vsx1*CreERT^2^ adult mouse retina suggests *Vsx1* expression in type 1 bipolar cells. Axon stratification of subsets of bipolar cells in S2 in postnatal day 3, 5, and 7 tamoxifen injected *Vsx1*CreER^T2^ mice also suggest possible temporal expression of *Vsx1* in type 3 bipolar cells during the first postnatal week. This is consistent with previous studies and suggests temporal *Vsx1* expression in non-canonical subtypes during bipolar cell development (Shekhar, et al. 2016; Shi, et al. 2012).

Balasubramanian, et al. (2014) showed *Lhx3* expression in ON and OFF cone bipolar cells and scRNA-Seq study by Shekhar, et al. (2016) later showed this expression specifically type 1b, 2, and 6 bipolar cells as confirmed in *Lhx3*CreER^T2^ adult mouse retinas in this study. *Lhx3*CreER^T2^ adult mouse retina had the highest percent BHLHB5+ type 2 bipolar cells and type 6 bipolar cells (the only ON bipolar cell) suggesting *Lhx3*’s role in the maintenance of these subtypes. *Lhx4* has also been shown to be expressed in a number of bipolar cell subtypes – specifically in types 2, 3, 4, and 5 bipolar cells and transiently in rod bipolar cells (Dong, et al. 2020). Shekhar, et al. (2016) also showed weak *Lhx4* expression in type 1a, 6, and rod bipolar cells in the postnatal day 17 mouse retina. Similar *Lhx4* expression pattern was observed when Cre was induced in the postnatal and adult *Lhx4*CreER^T2^ retina in our studies. *Lhx4* expression is detected in 8 out of 15 identified bipolar cell subtypes in the adult retina and thus the high density of bipolar cells in adult *Lhx4*CreER^T2^ retinas. Of note, *Lhx4* expression in the outer neuroblastic layer of the embryonic retina and inner nuclear layer of the postnatal retina (Blackshaw, et al. 2004) suggests its role in differentiation of all or a larger subset of bipolar cells the developing retina. This is also highlighted by *Lhx4*’s expression in atypical ON bipolar cell subtypes early in the first postnatal week and its later restriction to typical types 2-5 bipolar cell subtypes in the adult mouse retina. *Lhx4* expression in rod bipolar cells in the first postnatal week also appears to gradually decline as rod bipolar cells develop in the mature retina. These findings show the significance of differential expression of transcription factors in bipolar cell subtypes at different timepoints during the course of retinal development. It is plausible that a number of these transcription factors are expressed in specified bipolar cells and are differentially shutdown during subtype differentiation. *Lhx4* has been shown to be expressed in nascent cone photoreceptor cells and conditional knockout of *Lhx4* has resulted in defects in photoreceptor function (Buenaventura, et al. 2019; Dong, et al. 2020). Similarly, we detected *Lhx4* expression in photoreceptors in the postnatal retina and this expression was maintained in a subset of cone photoreceptors in the adult retina. This expression pattern of *Lhx4* enables the investigation of the structural and functional interaction of photoreceptor cells and bipolar cells using the *Lhx4*CreER^T2^ mouse. *Vsx1* expression is detected in V2 interneuron precursors in the developing spinal cord (Francius, et al. 2016), and *Lhx3* and *Lhx4* expression is also found in the developing pituitary gland, hindbrain, spinal cord, and lung (Li, et al. 1994; Weng, et al. 2006; Zhadanov, et al. 1995). Thus, these inducible Cre models can also be used to assess and genetically modulate mechanisms involved in the development of these tissues in mice. The inducible nature of these models can also be utilized to control the time and degree at which genetic manipulation is performed.

The study characterizes the expression of *Vsx1*, *Lhx3*, and *Lhx4* in the adult mouse retina in contrast to their expression in the postnatal retina during bipolar cell specification and subtype differentiation. The use of ChR2EYFP as a reporter of gene expression also highlights the subtype morphology of bipolar cells and can be utilized to assess the electrophysiological activity of bipolar cell subtypes in the presence and absence of light stimulus. The study, however, showed structural but not functional characteristics of bipolar cell subtypes and leaves room for characterization of the functional properties of these subtypes as well as signaling interactions between photoreceptor cells and bipolar cells in future studies.

Together, we characterize the efficiency of bipolar cell subtype – specific inducible Cre mouse lines that can be utilized for further study of the structural and functional development of bipolar cell subtypes in postnatal and adult mouse retinas. We hereby also show the differential transcription factor expression patterns that occur in bipolar cells during their development and likely regulate bipolar cell subtype differentiation and maintenance.

## Author Contributions

LG and EQ designed research; EQ and LK performed research; EQ analyzed data; EQ and LG wrote the paper; XX and JS contributed unpublished reagents/analytic tools

## Acknowledgements

Authors would like to thank the Augusta University Imaging Core for their support

## References

Balasubramanian, R., Bui, A., Ding, Q., Gan, L. (2014) Expression of LIM-homeodomain transcription factors in the developing and mature mouse retina. Gene Expression Patterns, 14:1–8.

Blackshaw, S., Harpavat, S., Trimarchi, J., Cai, L., Huang, H., Kuo, W.P., Weber, G., Lee, K., Fraioli, R.E., Cho, S.-H. (2004) Genomic analysis of mouse retinal development. PLoS biology, 2:e247.

Buenaventura, D.F., Corseri, A., Emerson, M.M. (2019) Identification of genes with enriched expression in early developing mouse cone photoreceptors. Investigative Ophthalmology & Visual Science, 60:2787–2799.

Burmeister, M., Novak, J., Liang, M.-Y., Basu, S., Ploder, L., Hawes, N.L., Vidgen, D., Hoover, F., Goldman, D., Kalnins, V.I. (1996) Ocular retardation mouse caused by Chx10 homeobox null allele: impaired retinal progenitor proliferation and bipolar cell differentiation. Nature genetics, 12:376–384.

Chow, R.L., Volgyi, B., Szilard, R.K., Ng, D., McKerlie, C., Bloomfield, S.A., Birch, D.G., McInnes, R.R. (2004) Control of late off-center cone bipolar cell differentiation and visual signaling by the homeobox gene Vsx1. Proceedings of the National Academy of Sciences, 101:1754–1759.

Daigle, T.L., Madisen, L., Hage, T.A., Valley, M.T., Knoblich, U., Larsen, R.S., Takeno, M.M., Huang, L., Gu, H., Larsen, R. (2018) A suite of transgenic driver and reporter mouse lines with enhanced brain-cell-type targeting and functionality. Cell, 174:465–480. e22.

Della Santina, L., Kuo, S.P., Yoshimatsu, T., Okawa, H., Suzuki, S.C., Hoon, M., Tsuboyama, K., Rieke, F., Wong, R.O. (2016) Glutamatergic monopolar interneurons provide a novel pathway of excitation in the mouse retina. Current Biology, 26:2070–2077.

Dong, X., Xie, X., Guo, L., Xu, J., Xu, M., Liang, G., Gan, L. (2019) Generation and characterization of Lhx4tdT reporter knock-in and Lhx4loxP conditional knockout mice. genesis, 57:e23328.

Dong, X., Yang, H., Zhou, X., Xie, X., Yu, D., Guo, L., Xu, M., Zhang, W., Liang, G., Gan, L. (2020) LIM-homeodomain transcription factor LHX4 is required for the differentiation of retinal rod bipolar cells and OFF-cone bipolar subtypes. Cell reports, 32.

Duan, X., Krishnaswamy, A., De la Huerta, I., Sanes, J.R. (2014) Type II cadherins guide assembly of a direction-selective retinal circuit. Cell, 158:793–807.

Feng, L., Xie, X., Joshi, P.S., Yang, Z., Shibasaki, K., Chow, R.L., Gan, L. (2006) Requirement for Bhlhb5 in the specification of amacrine and cone bipolar subtypes in mouse retina.

Francius, C., Hidalgo-Figueroa, M., Debrulle, S., Pelosi, B., Rucchin, V., Ronellenfitch, K., Panayiotou, E., Makrides, N., Misra, K., Harris, A. (2016) Vsx1 transiently defines an early intermediate V2 interneuron precursor compartment in the mouse developing spinal cord. Frontiers in molecular neuroscience, 9:145.

Ghosh, K.K., Bujan, S., Haverkamp, S., Feigenspan, A., Wässle, H. (2004) Types of bipolar cells in the mouse retina. Journal of Comparative Neurology, 469:70–82.

Guo, L., Xie, X., Wang, J., Xiao, H., Li, S., Xu, M., Quainoo, E., Koppaka, R., Zhuo, J., Smith, S.B. (2023) Inducible Rbpms-CreERT2 Mouse Line for Studying Gene Function in Retinal Ganglion Cell Physiology and Disease. Cells, 12:1951.

Hack, I., Frech, M., Dick, O., Peichl, L., Brandstätter, J.H. (2001) Heterogeneous distribution of AMPA glutamate receptor subunits at the photoreceptor synapses of rodent retina. European Journal of Neuroscience, 13:15–24.

Haverkamp, S., Grünert, U., Wässle, H. (2000) The cone pedicle, a complex synapse in the retina. Neuron, 27:85–95.

Haverkamp, S., Specht, D., Majumdar, S., Zaidi, N.F., Brandstätter, J.H., Wasco, W., Wässle, H., tom Dieck, S. (2008) Type 4 OFF cone bipolar cells of the mouse retina express calsenilin and contact cones as well as rods. Journal of Comparative Neurology, 507:1087–1101.

Haverkamp, S., Wässle, H. (2000) Immunocytochemical analysis of the mouse retina. Journal of Comparative Neurology, 424:1–23.

Kretzschmar, K., Watt, F.M. (2012) Lineage tracing. Cell, 148:33–45.

Li, H., Witte, D., Branford, W., Aronow, B., Weinstein, M., Kaur, S., Wert, S., Singh, G., Schreiner, C., Whitsett, J. (1994) Gsh-4 encodes a LIM-type homeodomain, is expressed in the developing central nervous system and is required for early postnatal survival. The EMBO journal, 13:2876–2885.

Madisen, L., Mao, T., Koch, H., Zhuo, J.-m., Berenyi, A., Fujisawa, S., Hsu, Y.-W.A., Garcia III, A.J., Gu, X., Zanella, S. (2012) A toolbox of Cre-dependent optogenetic transgenic mice for light-induced activation and silencing. Nature neuroscience, 15:793–802.

Masu, M., Iwakabe, H., Tagawa, Y., Miyoshi, T., Yamashita, M., Fukuda, Y., Sasaki, H., Hiroi, K., Nakamura, Y., Shigemoto, R. (1995) Specific deficit of the ON response in visual transmission by targeted disruption of the mGluR6 gene. Cell, 80:757–766.

Mataruga, A., Kremmer, E., Müller, F. (2007) Type 3a and type 3b OFF cone bipolar cells provide for the alternative rod pathway in the mouse retina. Journal of Comparative Neurology, 502:1123–1137.

Quainoo, E., Gan, L. (2025) Bipolar Cell Development.

Shekhar, K., Lapan, S.W., Whitney, I.E., Tran, N.M., Macosko, E.Z., Kowalczyk, M., Adiconis, X., Levin, J.Z., Nemesh, J., Goldman, M. (2016) Comprehensive classification of retinal bipolar neurons by single-cell transcriptomics. Cell, 166:1308–1323. e30.

Shi, Z., Jervis, D., Nickerson, P.E., Chow, R.L. (2012) Requirement for the paired-like homeodomain transcription factor VSX1 in type 3a mouse retinal bipolar cell terminal differentiation. Journal of Comparative Neurology, 520:117–129.

Shi, Z., Trenholm, S., Zhu, M., Buddingh, S., Star, E.N., Awatramani, G.B., Chow, R.L. (2011) Vsx1 regulates terminal differentiation of type 7 ON bipolar cells. Journal of Neuroscience, 31:13118–13127.

Tsukamoto, Y., Morigiwa, K., Ueda, M., Sterling, P. (2001) Microcircuits for night vision in mouse retina. Journal of Neuroscience, 21:8616–8623.

Tsukamoto, Y., Omi, N. (2014) Some OFF bipolar cell types make contact with both rods and cones in macaque and mouse retinas. Frontiers in neuroanatomy, 8:105.

Valny, M., Honsa, P., Kirdajova, D., Kamenik, Z., Anderova, M. (2016) Tamoxifen in the mouse brain: implications for fate-mapping studies using the tamoxifen-inducible Cre-loxP system. Frontiers in cellular neuroscience, 10:243.

Weng, T., Chen, Z., Jin, N., Gao, L., Liu, L. (2006) Gene expression profiling identifies regulatory pathways involved in the late stage of rat fetal lung development. American Journal of Physiology-Lung Cellular and Molecular Physiology, 291:L1027–L1037.

West, E.R., Cepko, C.L. (2022) Development and diversification of bipolar interneurons in the mammalian retina. Developmental biology, 481:30–42.

Young, R.W. (1985) Cell proliferation during postnatal development of the retina in the mouse. Developmental Brain Research, 21:229–239.

Zagozewski, J.L., Zhang, Q., Pinto, V.I., Wigle, J.T., Eisenstat, D.D. (2014) The role of homeobox genes in retinal development and disease. Developmental biology, 393:195–208.

Zhadanov, A.B., Bertuzzi, S., Taira, M., Dawid, I.B., Westphal, H. (1995) Expression pattern of the murine LIM class homeobox gene Lhx3 in subsets of neural and neuroendocrine tissues. Developmental Dynamics, 202:354–364.

Zhang, X.M., Ng, A.H.L., Tanner, J.A., Wu, W.T., Copeland, N.G., Jenkins, N.A., Huang, J.D. (2004) Highly restricted expression of Cre recombinase in cerebellar Purkinje cells. genesis, 40:45–51.

